# An elevated environmental temperature impairs accumulation of the pattern recognition receptor FLS2

**DOI:** 10.1101/2025.10.20.683271

**Authors:** Bryony C.I.C. Jacobs, Kyle W. Bender, Emma Six, Cyril Zipfel, Marc R. Knight

## Abstract

Pattern-triggered immunity (PTI) is initiated when plants detect pathogen-associated molecular patterns (PAMPs) through pattern-recognition receptors (PRRs). How moderate increases in temperature affect this plant immune signalling remains unclear. We explored this by using flg22 and the leucine-rich repeat receptor kinase (LRR-RK) FLS2 as a model receptor-ligand system and Ca^2+^ signaling as a representative PTI output. A pre-treatment at 28 °C significantly impaired the flg22-induced [Ca^2+^]_cyt_ influx, leading to a reduced expression of calcium-dependent defence genes, *ICS1* and *EDS1.* This effect correlated with a temperature-dependent reduction in FLS2 abundance. A qualitatively similar inhibition of these responses was observed when membrane fluidity was artificially increased using benzyl alcohol. This suggests that the effect of elevated temperature might act through changes in membrane properties. Artificially restoring FLS2 protein levels rescued flg22-dependent Ca^2+^ signalling and *ICS1* and *EDS1* expression in seedlings pre-treated at 28°C or with benzyl alcohol. Together, these findings indicate that increased membrane fluidity reduces FLS2 protein levels, thereby compromising Ca^2+^ signalling, and probably other, flg22-indcued responses. This highlights a potential mechanistic link between temperature perception, membrane fluidity, and FLS2-dependent calcium signalling, providing insight into how an increase in global temperatures may compromise plant immune responses in the future.

## INTRODUCTION

Plants are continually challenged by fluctuating, and often devastating, environmental conditions. To optimise growth and fitness, they must rapidly and precisely assess conditions to activate an appropriate change in internal physiological processes. Central to this ability are complex cell signalling networks that detect and relay environmental information and coordinate an informed integrated response in context with other factors such as biotic stresses.

The second messenger calcium is a key orchestrator of responses to environmental parameters, converting external stimuli into intracellular signals that can activate the appropriate downstream responses. The properties of cytosolic calcium responses have been found to be stimulus-specific (McAinsh & Hetherington, 1998). These signals possess characteristic and specific kinetics and so are termed “calcium signatures” (McAinsh & Pittman, 2009). Emerging evidence suggests these calcium signatures are decoded by calcium-binding proteins, including calmodulin, calmodulin-like proteins, calcium-dependent protein kinases and calcineurin B-like proteins, to effect specific appropriate downstream responses (Hashimoto & Kudla, 2011; Poovaiah & Du, 2018; Kudla *et al*., 2018). Despite this knowledge, an intriguing question remains: do different combinations of environmental factors modify a calcium signature to make it contextually appropriate?

In the context of plant defence against microbial pathogens, calcium signalling plays a pivotal role in patten-triggered immunity (PTI), which is initiated by the recognition of pathogen-associated molecular patterns (PAMPs) by plant pattern recognition receptors (PRRs) (Jones & Dangl, 2006). A paradigm example is the perception of the bacterial flagellin-derived peptide, flg22, by the plasma-membrane leucine-rich repeat receptor kinase FLAGELLIN SENSING 2 (FLS2) (Felix *et al*., 1999; Gómez-Gómez & Boller, 2000). The binding of flg22 to FLS2 causes a rapid increase in cytosolic calcium concentration [Ca^2+^]_cyt_, mostly through the influx of calcium from external stores (Jeworutzki *et al*., 2010). This elevated calcium signal initiates, amongst other events, the expression of defence genes including *ISOCHORISMATE SYNTHASE 1* (*ICS1*) and *ENHANCED DISEASE SUSCEPTIBILITY 1* (*EDS1*), which mediate salicylic acid-dependent immune responses (Lenzoni *et al*., 2018). Calcium-dependent regulation of *ICS1* and *EDS1* occurs through the calcium-calmodulin regulated transcription factors CAMTA3 and CBP60g, respectively (Wang *et al*., 2009b; Zhang *et al*., 2010).

Amongst emerging climate risks, an increase in ambient environmental temperatures has been identified as one of the main abiotic stresses plants need to adapt to (Bita & Gerats, 2013; Suzuki *et al*., 2014; Velásquez *et al*., 2018; Desaint *et al*., 2021). More specifically, increases in average ambient temperatures due to climate change has significantly affected plant-pathogen interactions globally (Chaloner *et al*., 2021). It is believed that these temperatures can heighten pathogenicity by increasing phytopathogen virulence, fitness and/or reproduction rate (Deutsch *et al*., 2008; Vaumourin & Laine, 2018). In plant defence responses, this trend has generally favoured the pathogens (Garrett *et al*., 2006; Desaint *et al*., 2021), leading to a greater incidence and intensity of disease in crops including coffee leaf rust (*Hemileia vastatrix*), potato blight (*Phytophthora infestans*), citrus canker (*Xanthomonas* spp.), and wheat rust (*Puccinia* spp.) (Singh *et al*., 2023; Angelotti *et al*., 2024). At the molecular level, an increased temperature seems to have a disputed impact on PTI. One study suggests that a short exposure to moderately high temperatures (<32°C) enhances PTI-specific transcriptomic changes in Arabidopsis (Cheng *et al*., 2013). Several other studies, however, have shown that PTI-dependent signalling, gene expression and hormone biosynthesis are compromised by an increase in ambient temperature (Wang *et al*., 2009a; Rasmussen *et al*., 2013; Huot *et al*., 2017; Janda *et al*., 2019; Kim *et al*., 2022). Though the impact of an increased ambient temperature on PTI has been studied extensively, this has yet to reveal its impact upon PTI-specific calcium signalling.

As described above, calcium signalling is known to play a central role in plant defence signalling. How an increase in environmental growth temperature may impact this, however, remains unknown. In this work, we use [Ca^2+^]_cyt_ measurements, as a key marker of PTI-specific signalling, to explore the impact of an increase in ambient temperature on flg22-dependent responses. We initially detected a reduction in the upstream calcium signalling response to flg22 following a pre-treatment at an increased temperature. Subsequently, we tested downstream Ca^2+^ signalling by measuring the expression of *ICS1* and *EDS1* and found it to be similarly compromised. Under this higher ambient temperature treatment, basal FLS2 protein levels were reduced, which, at least partly, explains the reduced calcium-dependent flg22 response measured following this treatment. Additionally, the calcium responses (upstream and downstream) could then be restored by artificially inducing FLS2 protein levels, further suggesting that reduced PTI in response to higher ambient temperature is due to a reduced availability of FLS2. We also suggest than an increase in temperature affects FLS2 accumulation, and likely PTI, via changes in membrane fluidity. This conclusion is based on the results which show that chemically altering membrane fluidity with benzyl alcohol (BA) similarly reduced upstream and downstream calcium signalling via a reduction in FLS2 protein levels. This inhibition of PTI by BA via reduced FLS2 levels could also be restored by the inducible expression of FLS2. Our findings therefore reinforce the negative effect of an increased temperature upon PTI-dependent calcium signalling and dissects a potential mechanism underpinning this effect.

## RESULTS

### An increase in environmental temperature reduces the upstream [Ca^2+^]_cyt_-dependent response to flg22

To assess the effect of an increase in environmental temperature upon plant defence responses, we incubated Arabidopsis seedlings (pMAQ2) at 28°C for 24 h. This moderate temperature increase (+8°C from normal growth temperature) was specifically selected based on research showing that it serves as an environmental signal, rather than a stress stimulus (Penfield, 2008; Liu *et al*., 2015). Following this incubation, the subsequent [Ca^2+^]_cyt_ and ROS responses to flg22 was measured. As shown in Figure 1, seedlings pre-treated at 28°C exhibited a clear reduction in the magnitude of the calcium (1a) and ROS (1b) response compared to those pre-treated at 20°C. This was evident in both the alteration of the flg22-induced calcium signature and the significantly decreased flg22-specific area under the curve (AUC).

**Figure 1.**
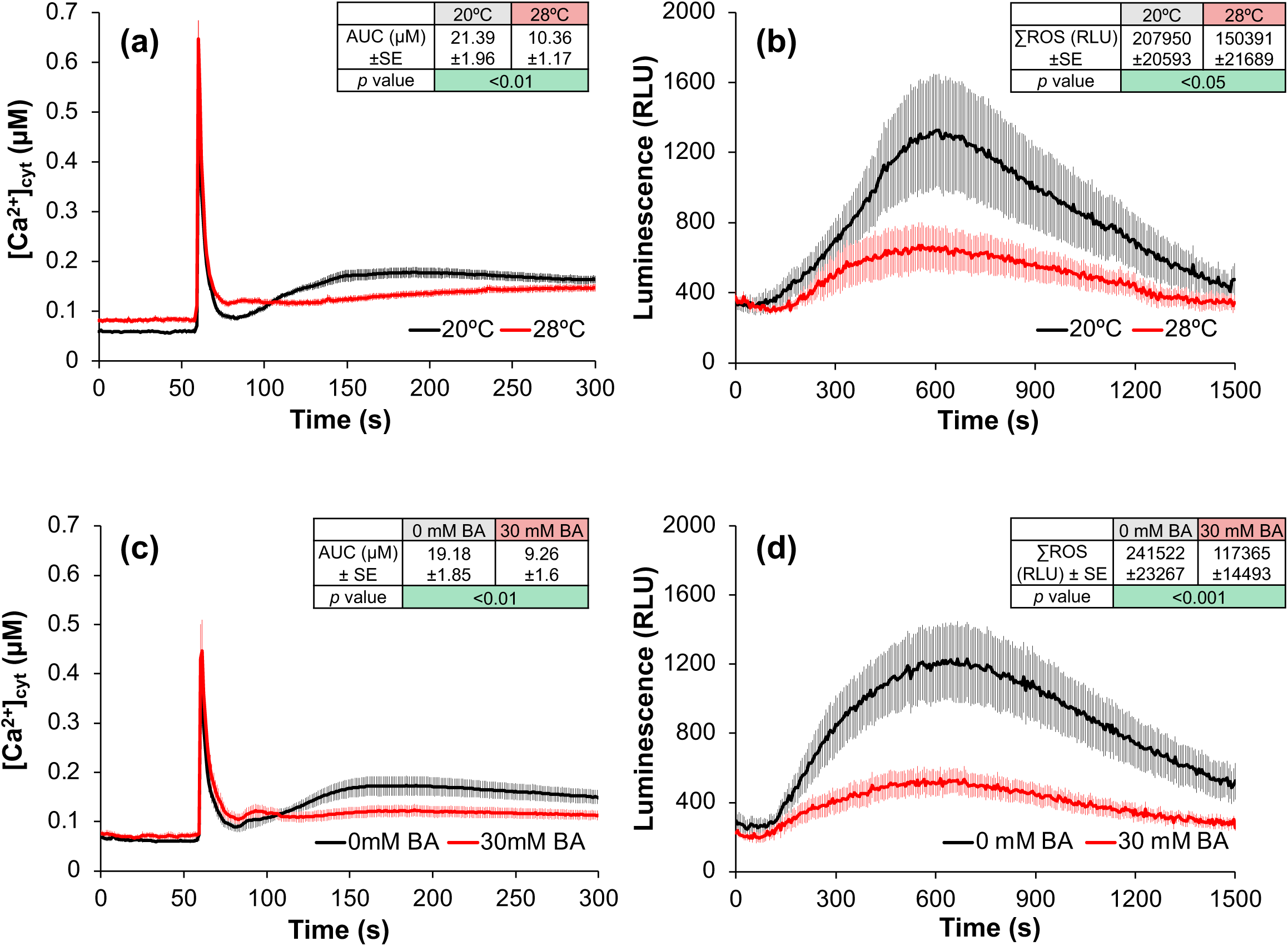
Flg22-induced [Ca^2+^]_cyt_ and ROS increases are quantitatively reduced in Arabidopsis seedlings pre-treated at 28°C or with benzyl alcohol (BA). [Ca^2+^]_cyt_ and ROS changes in response to 0.5 μM flg22 in Arabidopsis seedlings pre-treated for 24 h a) and b) at 20°C or 28°C or c) and d) at 20°C with 0mM (H_2_O) or 30mM BA. The traces shown are means of 15 replicate measurements and the error bars represent the S.E.M. For a) and c) the average area under the curve (AUC (µM)) ± S.E.M (SE) values for the flg22-induced calcium responses (100-300 s) are shown. The AUC values are the means of 15 replicate responses and the *p* value shown is the significance of the differences in the AUC as determined by a pairwise *t*-test. For b) and d) the average total ROS production (∑ROS (RLU)) ± S.E.M (SE) values for the flg22-induced ROS responses (0-1500 s) are shown. The ∑ROS values are the means of 15 replicate responses and the *p* value shown is the significance of the differences in the ∑ROS as determined by a pairwise *t*-test.

Many studies have shown that an elevated environmental temperature increases plant membrane fluidity (Murakami *et al*., 2000; Martinière *et al*., 2011; Cano-Ramirez *et al*., 2021). We therefore investigated whether the reduction in the flg22-dependent calcium response in seedlings incubated at 28°C was related to a temperature-induced increase in membrane fluidity. To test this, seedlings were treated with benzyl alcohol (BA), a compound known to artificially increase membrane fluidity, for 24 h (Carratù *et al*., 1996; Saidi *et al*., 2011; Niu & Xiang, 2018). As shown in Figures 1c and 1d, a BA pre-treatment, like a 28°C incubation, significantly reduced the [Ca^2+^]_cyt_ and ROS responses to flg22, respectively.

### An increase in environmental temperature also reduces the downstream [Ca^2+^]_cyt_-dependent response to flg22

Having shown that a 24 h incubation at 28°C reduces the early signalling responses to flg22 (Figure 1), we next tested whether this pre-treatment also suppressed downstream flg22-responsive calcium-dependent gene expression. *ICS1* and *EDS1* are two key biotic stress-inducible genes involved in the production of salicylic acid (SA) during plant defence responses. Both genes are also known to be regulated in a calcium-dependent manner by the transcription factors CBP60g and CAMTA3, respectively (Wang *et al*., 2009b; Zhang *et al*., 2010). Therefore, by using *ICS1* and *EDS1* expression as markers of downstream calcium-specific defence responses, we explored whether reducing the flg22-induced upstream signals with a 24 h incubation at 28°C (Figure 1), correlated with a reduced induction of these genes. Figure 2 supports this idea, with flg22-dependent *ICS1* and *EDS1* expression clearly reduced in the seedlings pre-treated at 28°C compared to those pretreated at 20°C.

**Figure 2.**
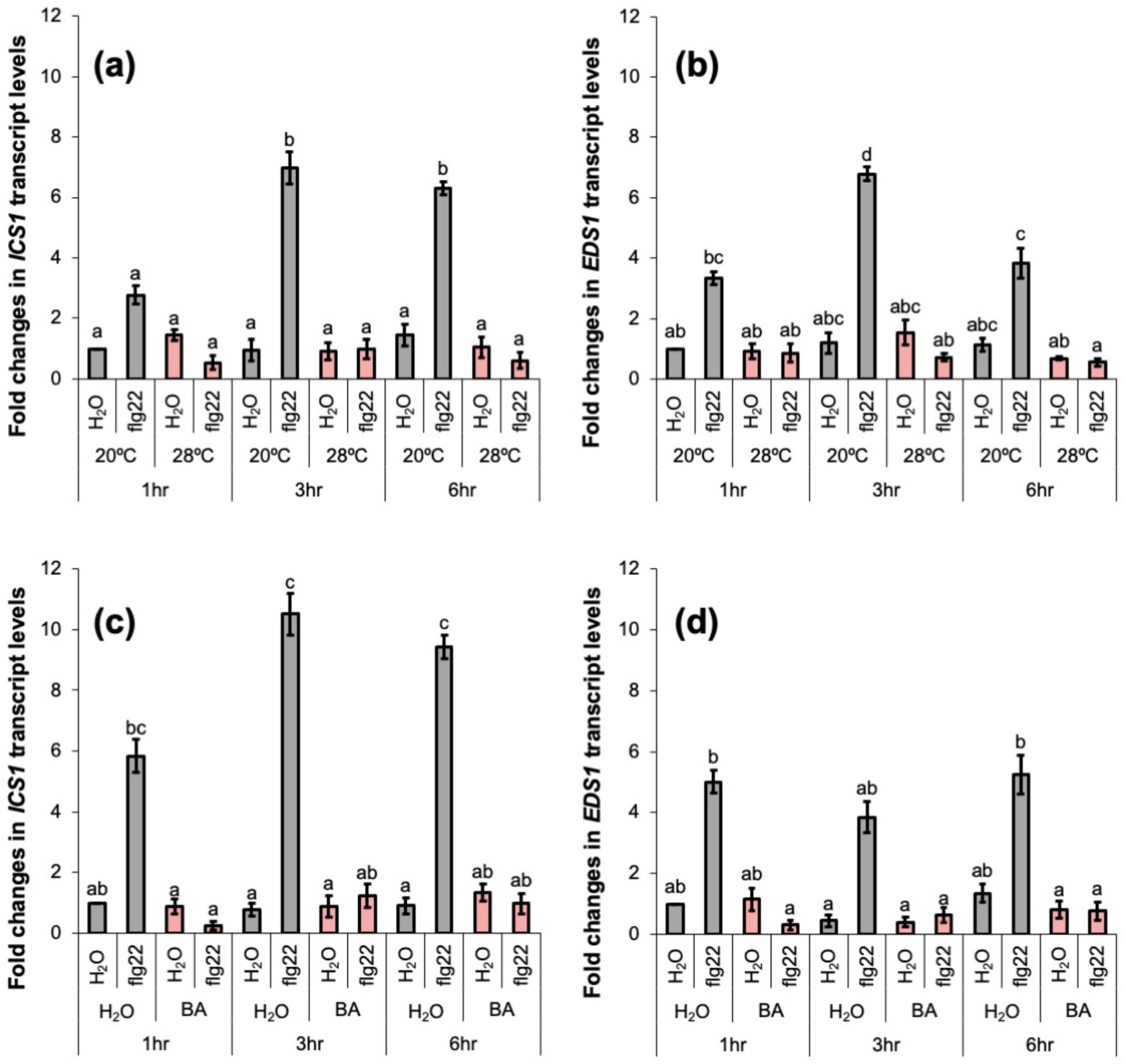
Flg22-dependent *ICS1* and *EDS1* expression is reduced in Arabidopsis seedlings pre-treated at 28°C or with benzyl alcohol (BA). Measurement by qPCR of the fold increases in a) and c) *ICS1* and b) and d) *EDS1* transcript expression in Arabidopsis seedlings pre-treated for 24 h at a) and b) 20°C or 28°C, or c) and d) at 20°C with 0 mM (H_2_O) or 30 mM benzyl alcohol (BA), in response to water or 0.5 μM flg22 1, 3, and 6 h after the start of treatment. Relative Quantification (RQ) values were calculated after normalisation to *PEX4* expression levels. The value produced for each treatment is the mean of three biological replicates, with each biological replicate representing the mean value of three technical repeats. Error bars represent the S.E.M and significant differences (*p* ≤0.05) were determined using ANOVA and the Tukey HSD post hoc test at a 95% confidence interval. Bars with the same letter are not significantly different.

To test whether the impact of temperature might be due to its effect upon membrane fluidity, we also measured the gene expression response to flg22 in seedlings pre-treated at 20°C with BA. We found that the BA pre-treatment shown to reduce the upstream [Ca^2+^]_cyt_ and ROS responses to flg22 (Figure 1), also led to a reduction in the flg22-dependent expression of *ICS1* and *EDS1* (Figure 2). The transcript level increase of both genes was significantly inhibited, at each time point, compared to the seedlings pre-treated with water. Together, these data demonstrate that reduction of the upstream calcium response to flg22, with either a 28°C or BA pre-treatment, correlates with reduced flg22-specific expression of calcium-regulated genes *ICS1* and *EDS1*.

### An increase in environmental temperature, or a treatment with BA, reduce *FLS2 levels*

The results presented thus far suggest that pre-treatment at an increased environmental temperature, or with BA, significantly reduces the upstream responses to flg22. To investigate the mechanism by which temperature/BA suppressed flg22 calcium-dependent responses, we next explored their effect on the level of the plasma membrane flg22 receptor, FLS2. Through western blot analysis we detected reduced basal FLS2 levels in seedlings pre-treated for 24 h at 28°C compared to those pre-treated at 20°C (Figure 3a). Similarly, BA pre-treatment also resulted in a clear comparative reduction in FLS2 protein levels (Figure 3b). With flg22-based signalling known to be dependent upon FLS2, a reduced level of FLS2 in seedlings pre-treated at 28°C and with BA (Figure 3) may explain why flg22-dependent upstream [Ca^2+^]_cyt_ signalling (Figure 1) and downstream gene expression (Figure 2) were also reduced in these seedlings.

**Figure 3.**
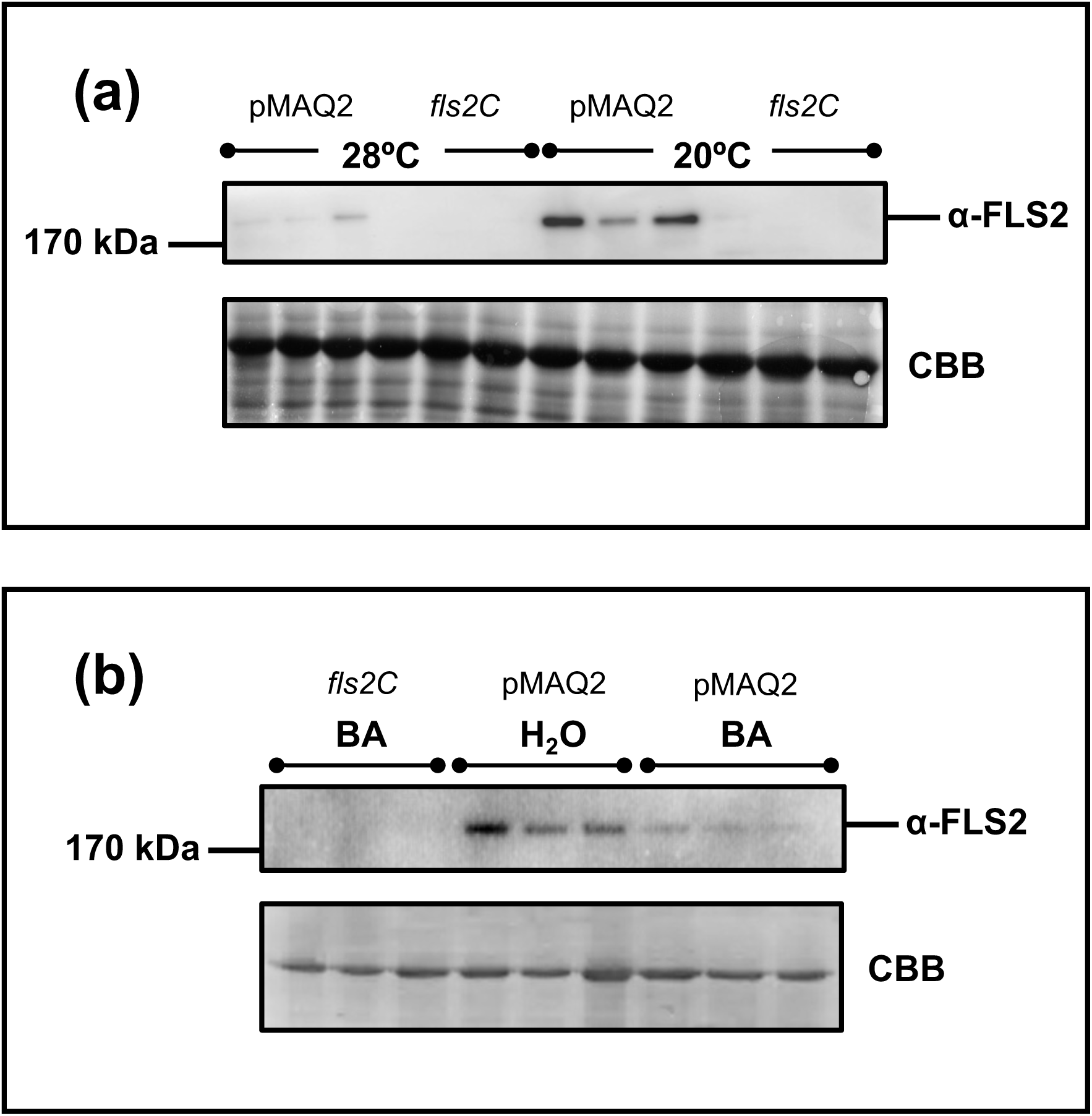
Basal FLS2 protein levels are lower in Arabidopsis seedlings pre-treated at 28°C or with benzyl alcohol (BA). Total protein was extracted from 14-day old pMAQ2 or *fls2C* (control) Arabidopsis seedlings pre-treated at a) 20°C or 28°C or b) at 20°C with 0 mM or 30 mM benzyl alcohol (BA) for 24 h. FLS2 levels were detected by western blotting. After detection, the blots were stained with Coomassie brilliant blue (CBB) to display loading. Values indicate the size (KDa) of the bands with the expected size of FLS2 indicated on the blot. Each sample loaded onto the blot is a biological replicate.

### FLS2 levels can be restored at 28*°*C upon inducible FLS2 over-expression

Our data so far suggest that a reduction in FLS2, induced by a pre-treatment at 28°C, contributes to the observed suppression in the calcium-dependent flg22 response. To manipulate basal FLS2 levels, we transformed pMDC7*FLS2*, which allows for *FLS2* transcription under an estradiol-inducible (XVE) system, into wildtype pMAQ2 plants. As a pre-treatment at 28°C (or with BA) was shown to reduce basal FLS2 levels (Figure 3), we firstly investigated whether the pMDC7*FLS2* construct could truly restore FLS2 protein levels in these conditions. Under control conditions (no estradiol), Figure 4 supports the previous findings: a reduced level of basal FLS2 is measured in wildtype seedlings pre-treated at 28°C or with BA, compared to those pre-treated at 20°C. More importantly, our data also show that FLS2 levels can be restored at 28°C, or with BA pre-treatment, following an estradiol treatment, as FLS2 protein can be clearly detected under these conditions. This suggests that we can use the pMDC7*FLS2* construct to artificially increase FLS2 levels in seedlings pre-treated at 28°C or with BA.

**Figure 4.**
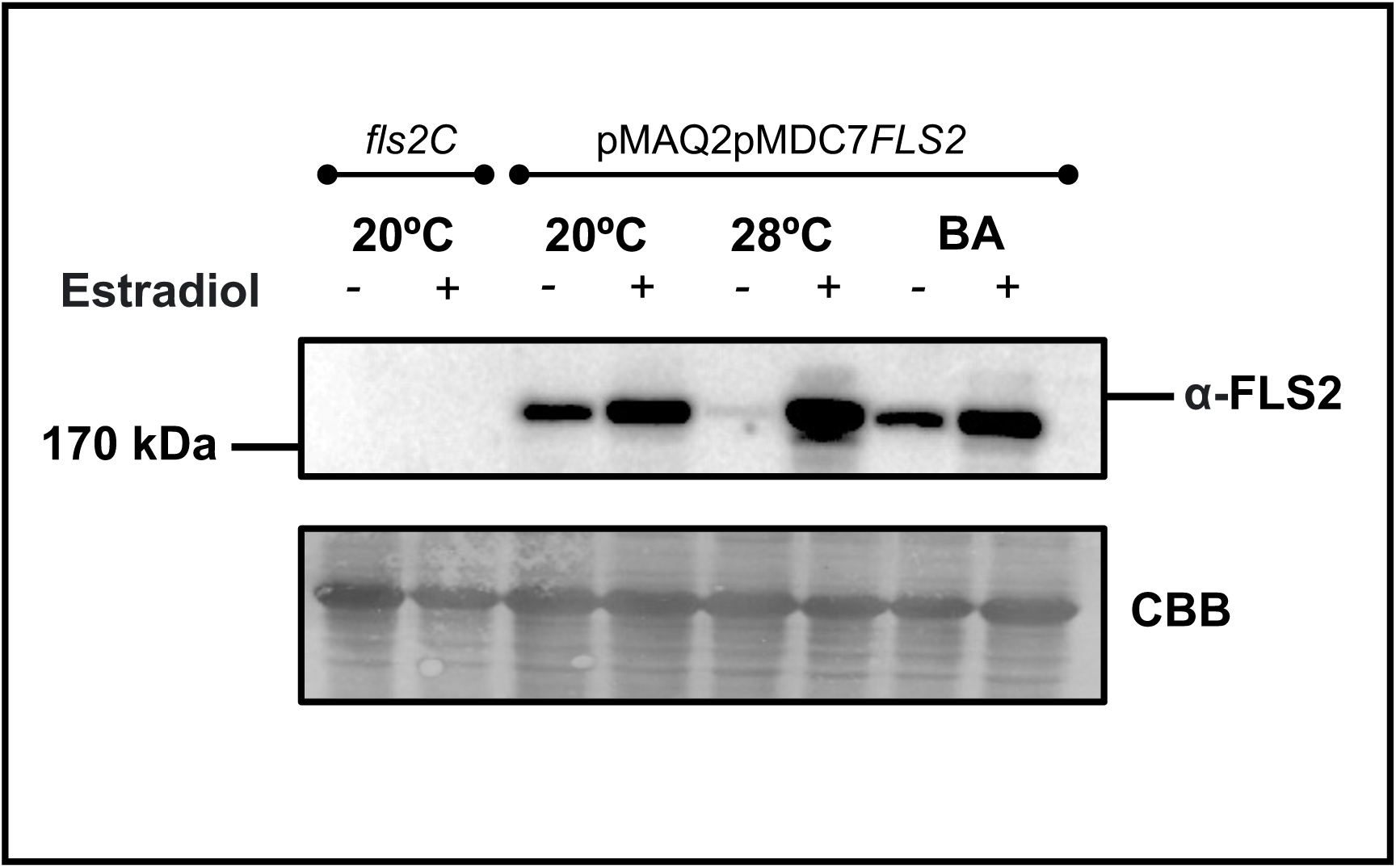
Basal FLS2 protein levels are restored in Arabidopsis seedlings pre-treated at 28°C or with benzyl alcohol (BA) using the pMDC7*FLS2* construct. Total protein was extracted from 14-day old pMAQ2pMDC7*FLS2* or *fls2C* (control) Arabidopsis seedlings pre-treated for 24 h at 20°C, 28°C or with 30 mM benzyl alcohol (BA). Seedlings also underwent a 16 h treatment of 10 μM estradiol (+) or 0.02% (v/v) DMSO (-) before protein extraction. FLS2 levels were detected by western blotting. After detection, the blots were stained with Coomassie brilliant blue (CBB) to display loading. Values indicate the size (KDa) of the bands with the expected size of FLS2 indicated on the blot.

### Restoring FLS2 levels at 28°C reinstates the calcium-dependent flg22 response at 28*°*C

After confirming the pMDC7*FLS2* construct restored FLS2 protein in wildtype seedlings pre-treated at 28°C, or with BA (Figure 4), we used the pMAQ2pMDC7*FLS2* line to investigate whether this restoration recovered flg22-dependent calcium signalling. To do this, we pre-treated seedlings for 24 h at 28°C or with 30 mM BA, together with a 16 h treatment of 10 μM estradiol or 0.02% (v/v) DMSO. The [Ca^2+^]_cyt_ response to flg22 was then tested in both sets of seedlings.

In the seedlings pre-treated with DMSO, we measured a partial restoration of the flg22-dependent response (Figure 5). This is seen in the characteristic flg22 [Ca^2+^]_cyt_ signature and an increased AUC measured in these seedlings compared to the pMAQ2 seedlings treated for 24 h at 28°C, or with BA, in Figure 1. This likely reflects some basal leakiness of the inducible system. Despite this, in the seedlings pre-treated with both 28°C and estradiol (Figure 5a), or both BA and estradiol (Figure 5b), an even larger increase in flg22-dependent calcium signalling was measured. The magnitude of the calcium signature and the total AUC were both significantly higher in these seedlings, suggesting a restoration (with estradiol) of FLS2 in pMAQ2pMDC7*FLS2* seedlings reinstates upstream flg22-specific calcium signalling following 28°C or BA pre-treatment

**Figure 5.**
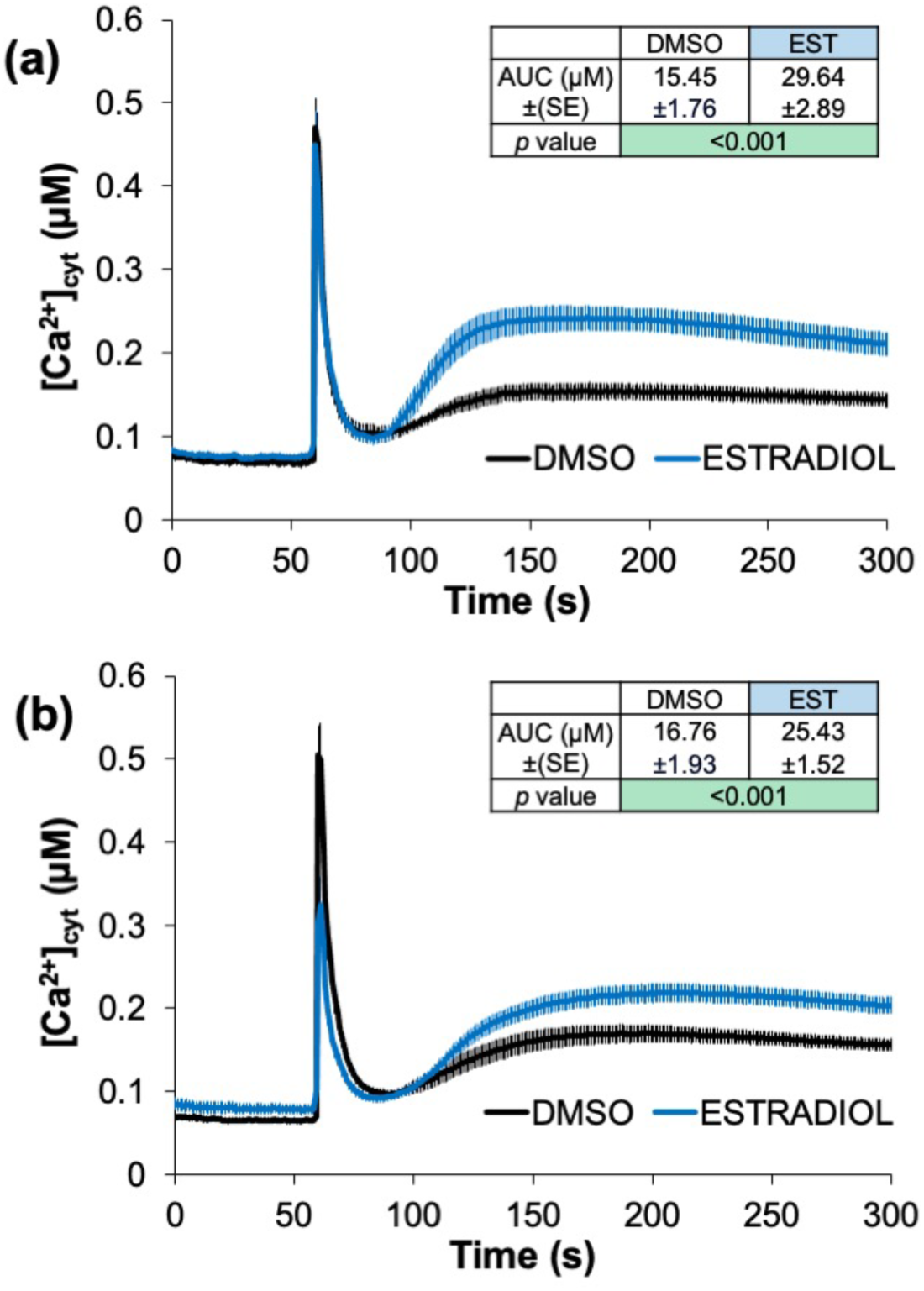
Flg22-induced [Ca^2+^]_cyt_ increases are restored in Arabidopsis seedlings pre-treated at 28°C or with benzyl alcohol (BA) using the pMDC7*FLS2* construct. [Ca^2+^]_cyt_ changes in response to 0.5 μM flg22 in pMAQ2pMDC7*FLS2* Arabidopsis seedlings pre-treated for 24 h a) 28°C or b) at 20°C with 30 mM BA and concurrently pre-treated for 16 h with 10 μM estradiol (EST) or 0.02% (v/v) DMSO. The traces shown are means of 15 replicate measurements and the error bars represent the S.E.M. The average area under the curve (AUC (µM)) ± S.E.M (SE) values for the flg22-induced calcium responses (100-300 s) are shown. The AUC values are the means of 15 replicate responses and the *p* value shown is the significance of the differences in the AUC as determined by a pairwise *t*-test.

To determine whether restoring the upstream [Ca^2+^]_cyt_ response also restored downstream signalling, we measured flg22-responsive calcium-dependent gene expression. To do this, we exposed pMAQ2pMDC7*FLS2* seedlings to pre-treatments of 20°C, 28°C or with BA at 20°C, together with treatments of DMSO or estradiol, and measured the *ICS1* and *EDS1* transcript level increases following a 3 h flg22 treatment. This timing was used as it was previously shown to be within the timeframe of optimal flg22-dependent gene expression for both *ICS1* and *EDS1* (Figure 2). As shown in Figure 6, a flg22-dependent increase in both *ICS1* and *EDS1* transcript level was measured in pMAQ2pMDC7*FLS2* seedlings pre-treated at 20°C and with DMSO. This was further increased in the seedlings pre-treated at 20°C and with estradiol. This suggests the increased artificial levels of FLS2 produced with the pMDC7*FLS2* construct not only enables the reinstatement of the calcium signature (Figure 5), but this signature is also functional in terms of regulating *ICS1* and *EDS1* transcription. Figure 6 also confirms previous results, that a BA or 28°C pre-treatment reduces calcium-specific flg22-responsive gene expression (see: DMSO H_2_O treatment). More importantly, we also show that using estradiol to initiate an increase in *FLS2* transcription in pMAQ2pMDC7*FLS2* seedlings pre-treated at 28°C, or with BA, restores flg22-specific *ICS1* and *EDS1* expression (Figure 6). Taken together, we show that pre-treating wildtype seedlings at 28°C or with BA significantly reduces the calcium-dependent response to flg22. This reduction, in both conditions, can be restored by using a pMDC7*FLS2* construct which artificially increases the levels of FLS2 in these seedlings. This in turn allows for the restoration of flg22-dependent upstream [Ca^2+^]_cyt_ signalling and downstream defense responsive gene expression.

**Figure 6.**
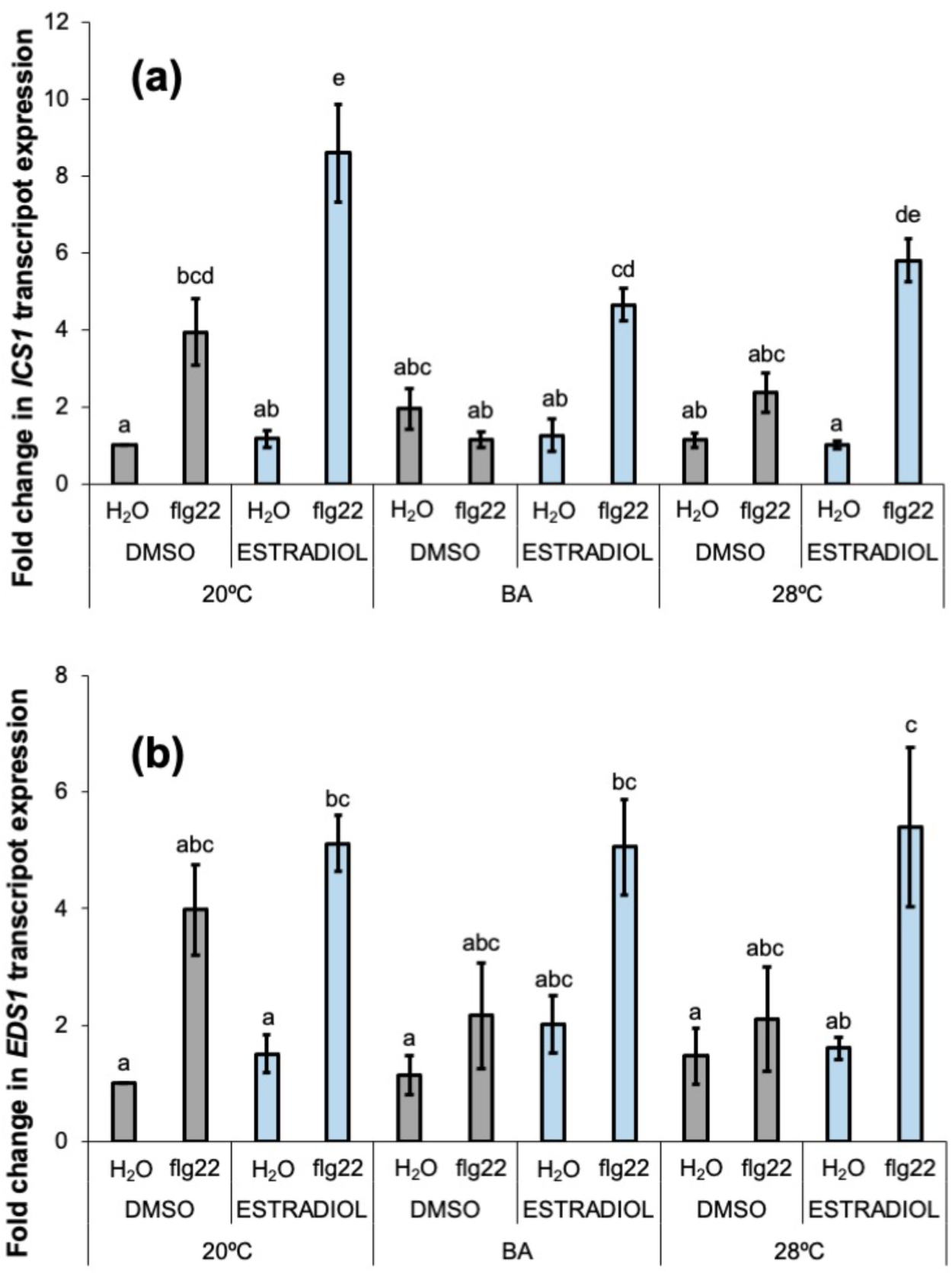
Flg22-dependent *ICS1* and *EDS1* expression is restored in Arabidopsis seedlings pre-treated at 28°C or with benzyl alcohol (BA) using the pMDC7*FLS2* construct. Measurement by qPCR of the fold increases in a) *ICS1* and b) *EDS1* transcript expression in pMAQ2pMDC7*FLS2* Arabidopsis seedlings pre-treated for 24 h at 20°C, 28°C or with 30 mM benzyl alcohol (BA) in response to water or 0.5 μM flg22 3 h after the start of treatment. The seedlings were also concurrently pre-treated for 16 h with 10 μM estradiol or 0.02% (v/v) DMSO. Relative Quantification (RQ) values were calculated after normalisation to *PEX4* expression levels. The value produced for each treatment is the mean of three biological replicates, with each biological replicate representing the mean value of three technical repeats. Error bars represent the S.E.M and significant differences (*p* ≤0.05) were determined using ANOVA and the Tukey HSD post hoc test at a 95% confidence interval. Bars with the same letter are not significantly different.

## DISCUSSION

The aim of this study was to investigate and examine the mechanistic basis of the impact of a moderate increase in temperature on the calcium-dependent signalling involved in Arabidopsis pattern-triggered immunity (PTI). To achieve this, we used the flg22-FLS2 ligand-receptor pair as a PTI model, and investigated [Ca^2+^]_cyt_ responses and quantified the expression of calcium-regulated immunity genes, *ICS1* and *EDS1* following a 24h pre-treatment at 28°C. We also measured basal FLS2 protein levels and tested the effect of modifying these levels by using an inducible FLS2 expression system. In parallel, we also investigated the effect of a BA treatment on the same markers of upstream and downstream calcium signalling, to test whether the effects of an increase in ambient temperature might be sensed through changes in membrane fluidity. Together, our work indicates that an increase in ambient temperature leads to desensitisation of PTI through the reduction in the amount of active FLS2, and that the increase in temperature is likely sensed by plant cells through an increased membrane fluidity.

Data in Figure 1a and 1b show that Arabidopsis seedlings exposed for 24 h at 28°C display significantly reduced upstream responses to flg22, compared to those kept at 20°C. This is seen as a clear statistically significant difference in the area under the curve for the later phase of the flg22-specific calcium signature and a reduced flg22-dependent ROS response. Very similar effects were observed when plants were pretreated for 24 h at 20°C with BA (Figure 1c and d). How the attenuation of upstream flg22-mediated responses correlate to the expression of flg22-induced calcium-regulated genes was then tested (Figure 2). For *EDS1* (Figures b and d) and *ICS1* (Figures a and c), both treatment at 28°C (Figures a and b) or with BA (Figures c and d) significantly inhibited their flg22-induced expression. This suggests a clear correlation between the [Ca^2+^]_cyt_ response and the expression of calcium-regulated genes *EDS1*/*ICS1*, and that both are reduced by a moderate increase in ambient temperature or with a treatment of BA. The similarity between the effects of 28°C and BA upon calcium signalling in response to flg22 suggests that the sensing of temperature increase that leads to a reduction in sensitivity of PTI acts through plant cells assessing changes in membrane fluidity. It is known that cells remodel membrane fluidity in response to changes in ambient temperature through controlling lipid saturation and fatty acid length (Schroda *et al*., 2015). Literature provides a consensus that this change in the physical state of the membrane can act as a “thermometer” (Niu & Xiang, 2018; Cano-Ramirez *et al*., 2021; Jung *et al*., 2023), whereby the biophysical increase in membrane fluidity caused by increases in temperature is the parameter sensed by cells to alert them of temperature change. Our experiments using BA, which will fluidise the membrane whilst keeping a constant temperature (Pedersen & Cox, 1984; Örvar *et al*., 2000; Sangwan *et al*., 2001), phenocopied the effects of 28°C treatment on both cytosolic calcium responses and downstream plant immunity gene expression. This is consistent with the hypothesis that perception of increased ambient temperature leading to reduction of FLS2 levels occurs via sensing the changes in membrane fluidity.

Precisely how this disruption in membrane fluidity is sensed and then relayed to effect flg22-dependent Ca^2+^ signal is an interesting conundrum. The increase in membrane fluidity may act as a signal itself, detected within the plasma membrane. Although several ion channels, notably the transient receptor potential cation channels (TRPs), have been shown to function as temperature sensors in animal cells (Caterina *et al*., 1997; Xu *et al*., 2002; Vilar *et al*., 2020), the identity of plant temperature sensors remains largely elusive. Specific calcium-permeable cyclic nucleotide gated channels (CNGCs) have been shown to be activated by heat and mild temperature increments in plants (Saidi *et al*., 2009; Finka *et al*., 2012; Gao *et al*., 2012). It may be interesting to determine the role of such channels in our work, especially as specific plasma membrane bound Ca^2+^ channels were shown to be activated, and modulated, by an increase in temperature or by chemically perturbating membrane fluidity with BA (Saidi *et al*., 2010). It is important to remember that the alteration in membrane fluidity serves also a physical cue, that may impact the conformation and function of plasma membrane-bound proteins and channels. Rather than a signal *per se*, it could be that the disruption in membrane fluidity inhibits the environment of important receptors, including FLS2. For example, with several studies concluding that FLS2 exists within PM nanoclusters, it could be important to measure the effect of changes in membrane fluidity upon nanocluster integrity (Bücherl *et al*., 2017; Cui *et al*., 2018; Tran *et al*., 2020; Gronnier *et al*., 2022; Hurst *et al*., 2023). This aligns with research showing that altering membrane composition and plasma membrane sterol abundance affects FLS2-based signalling (Cui *et al*., 2018; Hurst *et al*., 2023). Deciphering how the temperature-dependent impact on membrane fluidity is relayed to reduce Ca^2+^-specific PTI signalling would reveal further insight into the mechanism behind our work.

Though the source/nature of the signal from the membrane is currently unknown, it is still very clear that an increased ambient temperature or BA treatment reduce both upstream and downstream calcium signalling to flg22. As FLS2 is the flg22 receptor, we investigated the impact an increase in membrane fluidity has on its basal level. The specific interaction between flg22 and FLS2 has been shown to be responsible for initiating the calcium influx from external stores which creates the calcium signature we observed in Figure 1 (Jeworutzki *et al*., 2010; Ranf *et al*., 2011). To investigate whether an increase in ambient temperature and/or BA might affect the levels of this receptor, we performed western blot analysis on seedlings pre-treated for 24 h at either 28°C (Figure 3a) or with BA (Figure 3b). This analysis clearly showed that total levels of FLS2 protein were reduced (Figure 3) in these conditions. This suggested that the reduction in level of upstream (Figure 1) and downstream (Figure 2) signalling in response to flg22 that we observed was due to a reduction in the level of the primary receptor. To test this hypothesis, we generated a DNA construct which, when transformed into Arabidopsis, could induce FLS2 expression upon exogenous application of estradiol. Expression of this construct was used to see whether by judiciously inducing expression of FLS2 under conditions (28°C or BA) which led to a reduction in total FLS2 protein seen in Figure 3 could restore calcium signalling in response to flg22. We first tested whether this construct could achieve increased FLS2 protein levels under these conditions. As can be seen in Figure 4, under conditions of 28°C or BA, FLS2 protein levels were reduced as already observed (Figure 3), but estradiol treatment led to restoration of FLS protein levels. As can be seen in Figure 5, when FLS2 was specifically expressed after 24 h of 28°C or BA at 20°C, this restored the calcium response to flg22, with a significantly increased area under the curve being produced. To test whether this restoration of flg22-mediated upstream calcium response could also restore downstream *ICS1* and *EDS1* expression, we tested the same conditions and measured transcript levels of these 2 genes. As can be seen in Figure 6 whilst 28°C and BA treatments again inhibited the flg22-mediated induction of both genes, compared to expression at 20°C, restoration of FLS2 by estradiol treatment restored the ability of these plants to respond to flg22. These data together strongly support the idea that the desensitisation of the response at both 28°C and in response to BA treatment is due to a reduction in active FLS2 receptor levels. It has been shown previously that an acute (45 minutes) treatment at high temperature (42°C) greatly reduces the ROS response to flg22 due to reduced FLS2 (Janda *et al*., 2019). Our work shows that similar effects can be seen in response to much more moderate temperature elevations, which are much closer to relevant field temperatures, suggesting this phenomenon is of significant contemporary relevance to agriculture. The impact of a smaller temperature increase upon FLS2 we measured in this work may help to explain some of the well-established research showing suppression of PTI-based signalling under more moderate temperature increases. For example, FLS2-dependent callose deposition (Gómez-Gómez & Boller, 2000; Zipfel *et al*., 2004) in response to flg22 is reduced in plants exposed to a 24 h pre-treatment at 37°C (Janda *et al*., 2019) or 48 h pre-treatment at 30°C (Huot *et al*., 2017). Even more related to our work, transcriptomic analysis has previously shown a reduction in flg22-dependent *ICS1* and *EDS1* gene expression following a short pre-treatment at 37°C (Rasmussen *et al*., 2013). This suppression of pathogen-induced *ICS1* expression has also been measured following both a 30°C (Huot et al., 2017), and 28°C (Shields *et al*., 2025) treatment, which correlates with the subsequent reduction of salicylic acid production at 28°C compared to 23°C (Kim *et al*., 2022). This work clearly aligns with the data in our study, suggesting the suppression of FLS2 at 28°C may be involved in a more global suppression of flg22-dependent responses following increased ambient temperatures. Though it is clear the temperature-specific reduction of FLS2 strongly impairs PTI signalling, in the future we will investigate the mechanism by which FLS2 levels are reduced in response to temperature and BA. It is most likely that these treatments impose destabilisation of the protein and protein degradation, therefore measuring ubiquitination of FLS2 in response to 28°C/BA would be a good approach. Equally, it is possible that mechanisms that have been described for autophagic regulation of FLS2 could be involved (Yang *et al*., 2019). Increases in ambient temperature are well-known inducers of autophagy (Sedaghatmehr *et al*., 2019) and so it would be interesting if the ORM1/ORM2 orosomucoid proteins which selectively degrade FLS2 through ATG8 autophagy (Yang *et al*., 2019), are increased in expression or activity in response to increased ambient temperature.

Overall, the work described here demonstrates that a potential component in the increase in balance of power in favour of pathogens of crops due to modest increases in temperature during climate change could be due to destabilisation of receptors evolved to detect pathogens. Understanding this, and future research into the mechanisms by which these receptors are regulated by increases in temperature might be useful for reading/engineering crops with robustness of receptor levels under increasing temperature. Targets could be the components of the ubiquitin pathway specifically regulating this phenomenon, the temperature-dependent autophagy pathway leading to FLS2 or membrane micro domain and lipid species that govern FLS2 stability at elevated temperature. In addition, it will be interesting to expand this work to investigate whether the reduction in FLS2 levels is seen similarly in other PRRs.

## MATERIAL AND METHODS

### Plant materials and growth conditions

All seedlings used were in the *Arabidopsis thaliana* (*A. thaliana*) Col-0 ecotype background. Wildtype calcium measurement experiments were performed on transgenic seedlings constitutively expressing the calcium reporter 35S::apoaequorin in the cytosol (Col-0pMAQ2) (Knight *et al*., 1991). The *fls2-26*pMAQ2 mutant, which constitutively expresses cytosolic 35S::apoaequorin, and possesses a nucleotide missense mutation (Q865*) in *FLS2*, was described previously (Ranf *et al*., 2012). The *fls2C* mutant (SAIL_691C4) was used as a *FLS2* null mutant in the immunoblotting work (Zipfel *et al*., 2004). The pMAQ2pMDC7*FLS2* line used was created in this work. Seeds were surface sterilised in 70% ethanol (v/v) and sown onto 1 × Murashige and Skoog (MS; Duchefa Biochemie BV, Haarlem, Netherlands) medium, pH 5.8, 0.8% (w/v) plant tissue culture agar (Sigma-Aldrich, St Louis, MO, USA) and stratified at 4°C in darkness for 48 h. Plants were then grown in a Percival CU-36L5D growth chamber (CLF PlantClimatics, Emersacker, Germany) at 20±1°C, with a light intensity of 150 μmol m^-2^ s^-1^ and a 16 h light/8 h dark photoperiod. Pre-treatments were conducted 24 h before the start of experiments with specific details given below. All experiments were performed on 14-day-old seedlings.

### Producing the pMDC7FLS2pMAQ2 line

The pENTR^TM^/D-TOPO^TM^ entry vector (Thermo Fisher Scientific, Loughborough, UK) was linearised using NotI and AscI restriction enzymes (New England Biolabs, UK), and the resulting product isolated from an agarose gel by gel extraction (QIAGEN Ltd., UK). Three synthetic gene blocks, designed to span across the full *FLS2* coding sequence (CDS), were ordered from IDT (Integrated DNA Technologies, Leuven, Belgium). The sequences of these gene blocks can be found in Supplementary Table 1. A Gibson Assembly® Cloning Kit (NEB, Cat. No E5510S) was used to assemble the *FLS2*CDS gene fragments into the linearised vector according to the manufacturer’s instructions. The binary destination vector pMDC7, containing the estradiol inducible XVE system, was described previously (Curtis & Grossniklaus, 2003). Gateway recombination using the LR Clonase^TM^ II Enzyme Mix (Life Technologies, Paisley, UK) was then performed between the pENTR*FLS2*CDS entry clone and the pMDC7 destination vector.

### In vivo reconstitution of aequorin and pre-treatment conditions

For calcium measurements, 24 h before the start of measurements 13-day-old seedlings were floated on a 5 mL H_2_O solution in 6-well plates. For temperature pre-treatment conditions, the seedlings were moved into one of two identical Sanyo MLR-351 growth cabinets (Sanyo Electric co. Ltd, Moriguchi, Japan). One growth cabinet was set at 20°C, and the other was set at 28°C. For benzyl alcohol (BA) pre-treatment conditions, seedlings were floated in either 5 mL H_2_O or 30 mM BA and placed inside the 20°C growth cabinet. Aequorin reconstitution was performed during these pre-treatment conditions by adding coelenterazine (final concentration 10 μM in 1% (v/v MeOH) to the solution 16 h before the start of experiments. Where used, estradiol pre-treatment was also conducted during this period by adding estradiol (final concentration: 10 μM in 0.02% (v/v) DMSO) or DMSO (final concentration 0.02%(v/v)) as a control to the solution 16 h before the start of the experiment.

### [Ca^2+^]_cyt_-dependent luminescence measurements

Following pre-treatment, individual seedlings were transferred into 3.5 mL luminometer cuvettes (Sarstedt, Nümbrecht, Germany) containing 0.5 mL H_2_O. After a 30 min period of rest, individual cuvettes were then inserted into the luminometer sample housing. Luminescence levels were recorded every 1 s using a digital chemiluminometer with discriminator and cooled housing unit (Electron Tubes Limited, Middlesex, UK) to reduce background noise (Knight *et al*., 1991). Luminescence was recorded for 60 s before the injection of 0.5 mL 1 μM flg22 (QRLSTGSRINSAKDDAAGLQIA) (GenScript Biotech, New Jersey, USA). The subsequent changes in luminescence were recorded for a further 240 s. A 300 s discharge was performed at the end of the experiment by injecting equal (1 mL) volume of 2M CaCl_2_, 20% (v/v) ethanol. Calibration was performed as described previously (Knight *et al*., 1991).

### ROS burst measurements

ROS burst measurements were performed on whole seedlings using an adapted version of a previously described protocol (Kadota *et al*., 2014). Briefly, individual seedlings were incubated overnight in luminometer cuvettes containing 17 μg/mL luminol (Sigma-Aldrich) and 20 μg/mL horseradish peroxidase (HRP, Sigma-Aldrich). The following day, the solution was replaced with a 0.5 μM flg22 solution (in 17 μg/mL luminol, 20 μg/mL HRP) and the individual cuvettes were then inserted into the luminometer sample housing. Luminescence levels were recorded every 1 s using a digital chemiluminometer with discriminator and cooled housing unit (Electron Tubes Limited, Middlesex, UK) to reduce background noise (Knight *et al*., 1991). Luminescence levels were recorded for 1500 s and ROS production is displayed as total luminescence recorded.

### RNA extraction, cDNA preparation and gene expression measurements

Pre-treatments were performed as described above for gene expression experiments. For treatments, 1 mL of solution was removed from each condition and 1 mL flg22 (or H_2_O) at 5 × concentration was added (in either 0 or 150 mM BA) and the seedlings were returned to either the 20°C or 28°C growth cabinet until harvested. Tissue was harvested 1, 3 and 6 h after the treatment. For each sample, representing one condition at a single time point, 15 seedlings were pooled together for subsequent RNA extraction. A high-capacity cDNA reverse transcriptase kit (Applied Biosystems, Foster City, CA, USA) was used to reverse transcribe total RNA (2 μg) obtained with an RNeasy ReliaPrep™ RNA Miniprep System Plant Total RNA kit (Promega, Southampton, UK). Quantitative real-time PCR was performed using 5 μL of 1:50 diluted cDNA in a total volume of 15 μL using an Applied Biosystem 7300 real time PCR machine. Relative expression of *Enhanced Disease Susceptibility 1* (*EDS1*) (*At3g48090*) and *Isochorismate Synthase 1* (*ICS1*) (*At1g74710*) were measured with Fast Start SYBR Green Master Mix with ROX (Promega, Southampton, UK) using the following primers: *EDS1* Fw 5′-ACCTAACCGAGCGCTATCAC-3′, *EDS1* Rev 5′-TTGTCCGGATCGAAGAAATC-3′, *ICS1* Fw 5′-CAAATCTCAACCTCCGTCGT-3′, *ICS1* Rev 5′-AATCAATTGCTCCGATTTGC-3′. Levels were normalised to the levels of the endogenous *PEX4* housekeeping gene (*At5g25760*), using the following primers: *PEX4* Fw 5′-TCATAGCATTGATGGCTCATCCT-3′ and *PEX4* Rev 5′-ACCCTCTCACATCACCAGATCTTAG-3′. Relative quantification was performed using the delta cycle threshold (ΔΔ*C_t_*) method (Livak & Schmittgen, 2001) and the values obtained representing the relative quantification (RQ) were calculated as described previously (Knight *et al*., 2009).

### Generation of anti-FLS2 antibodies

Anti-FLS2 antibodies were generated against a peptide targeting the C-terminus of FLS2 (CKANSFREDRNEDREV) and coupled to keyhole limpet hemocyanin for immunisation. Peptide synthesis, conjugation, and immunisations were performed by GenScript. Rabbit immune serum was used for affinity purification against bead-coupled antigen peptide. The specificity of affinity purified antibody was confirmed by immunoblotting against protein extracts from Arabidopsis seedlings lacking FLS2 protein (Supplementary Figure 1).

### SDS-PAGE and western blotting

Pre-treatments were performed as described above and samples were harvested directly after the 24 h pre-treatment (no flg22 treatment). For each sample, representing one condition, 15 seedlings were pooled together for subsequent protein extraction. Seedlings were flash frozen in liquid nitrogen and thoroughly homogenised in equal volume of extraction buffer (50 mM Tris-HCl pH 7.5, 150 mM NaCl, 10% (v/v) glycerol, 2 mM EDTA), 0.002M DTT, 1% (v/v) Igepal, and 1 × Phosphatase (P5726) and 1 × Protease (P9599) Inhibitors, Sigma-Aldrich). The samples were left on ice for 30 min to solubilise membrane proteins before being centrifuged at 20,000*g* for 20 min at 4°C. Each resulting 100 μL sample was normalised to contain the same amount of protein, and 35 μL 5 × SDS-PAGE loading buffer (10% SDS (w/v), 50% glycerol, 300 mM Tris-HCl pH 6.8, 0.125% (w/v) bromophenol blue) and 15 μL 1M DTT was added to each. Protein samples were heated to 90°C for 10 min prior to electrophoresis.

Samples were loaded onto 8% SDS-PAGE gels and electrophoresis was performed (in 25 mM Tris, 192 mM glycine, 0.1% SDS) at 150 V for approximately 2 h. Subsequently, proteins were transferred at 4°C onto activated PVDF membranes using wet transfer (in 25 mM Tris, 192 mM glycine, 20% (v/v) MeOH) at 30 V for 90 min. Membranes were subsequently blocked overnight with 5% (w/v) skimmed milk powder dissolved in fresh Tris buffered saline containing Tween-20 (TBS-T; 10 mM Tris-HCl pH 8, 150 mM NaCl, 0.1% (v/v) Tween-20) at 4°C with gentle (40 RPM) agitation. The primary antibody (α-FLS2, KLH-conjugated) was then added at a 1:2,500 dilution in TBS-T (5% (w/v) skimmed milk) to the membrane. The membrane was then incubated for 2 h at room temperature with gentle (40 RPM) agitation. After primary antibody incubation the membrane was washed five times with fresh TBS-T for 10 min each. The secondary antibody (goat anti-mouse IgG (H/L) HRP polyclonal antibody, Bio-Rad, USA) was added at a 1:5,000 dilution in TBS-T (5% (w/v) skimmed milk) to the membrane. Incubation occurred at room temperature with gentle (40 RPM) agitation for 1 h and the membrane was washed again with fresh TBS-T before detection.

Western blots were visualised using the SuperSignalTM West Femto Detection Kit (Thermo Fisher Scientific, Loughborough, UK) according to the manufacturer’s instructions. The substrate was distributed equally over the membrane and the HRP-dependent chemiluminescence was detected using either a ChemiDoc Imaging System (Bio-Rad, California, USA) or a photon counting camera.

## AUTHOR CONTRIBUTIONS

BCICJ and MRK designed the research, BCICJ, KWB and ES performed the experimental work, BCICJ performed the data analyses, collection and interpretation and BCICJ, MRK, KWB and CZ contributed to writing.

## ACKNOWLEDGEMENTS

This research was funded by the Biotechnology and Biological Sciences Research Council (BBSRC) through the award of a DTP PhD studentship (ref 2182091) to BCICJ, and core funding to CZ provided by the University of Zurich and the Gatsby Charitable Foundation. We would like to thank Stefanie Ranf for providing *fls2-26*pMAQ2 and Ueli Grossniklaus for the pMDC7 binary vector. We also acknowledge and thank the help given by Julia Davies.

---

**Supplementary Table 1.**
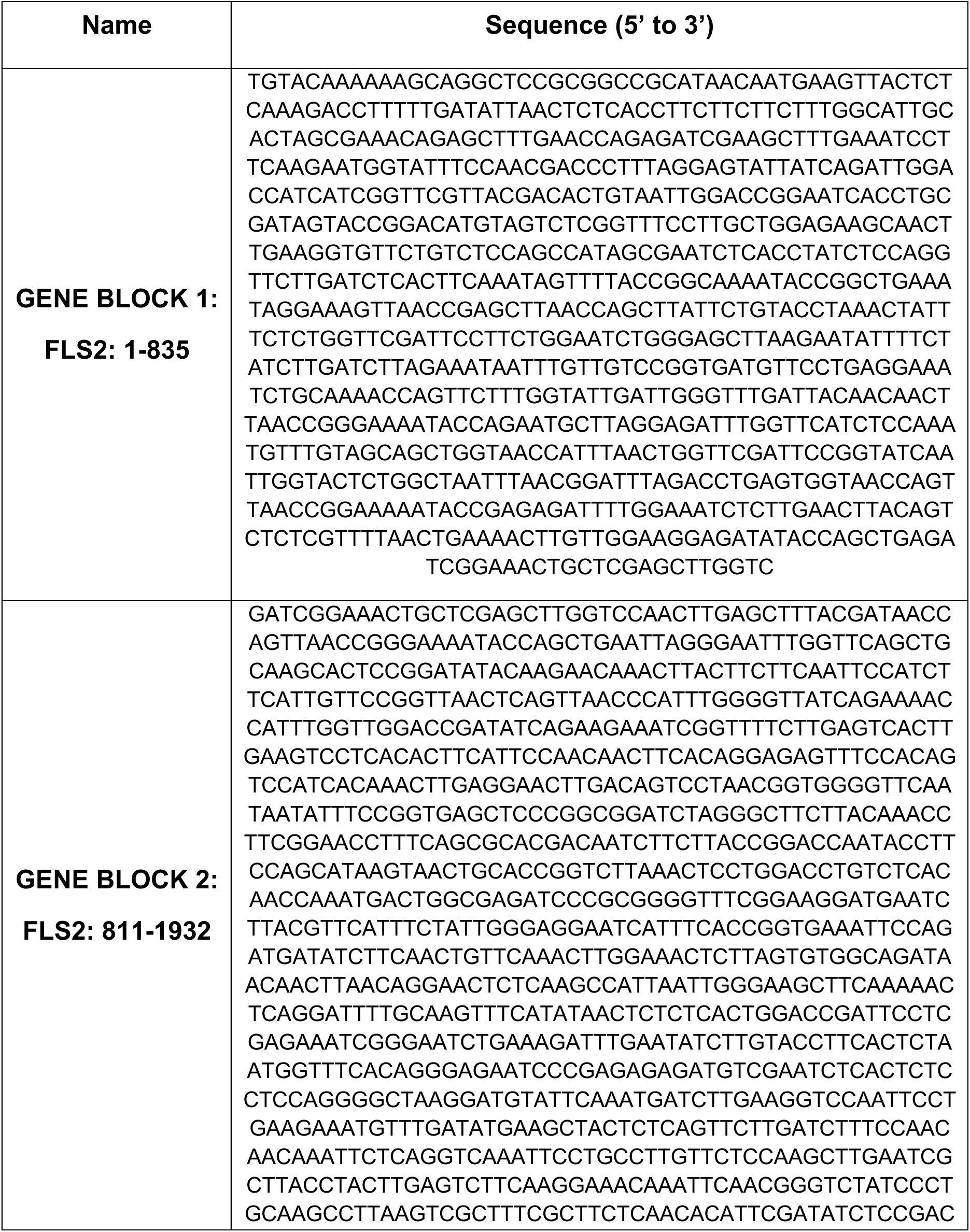
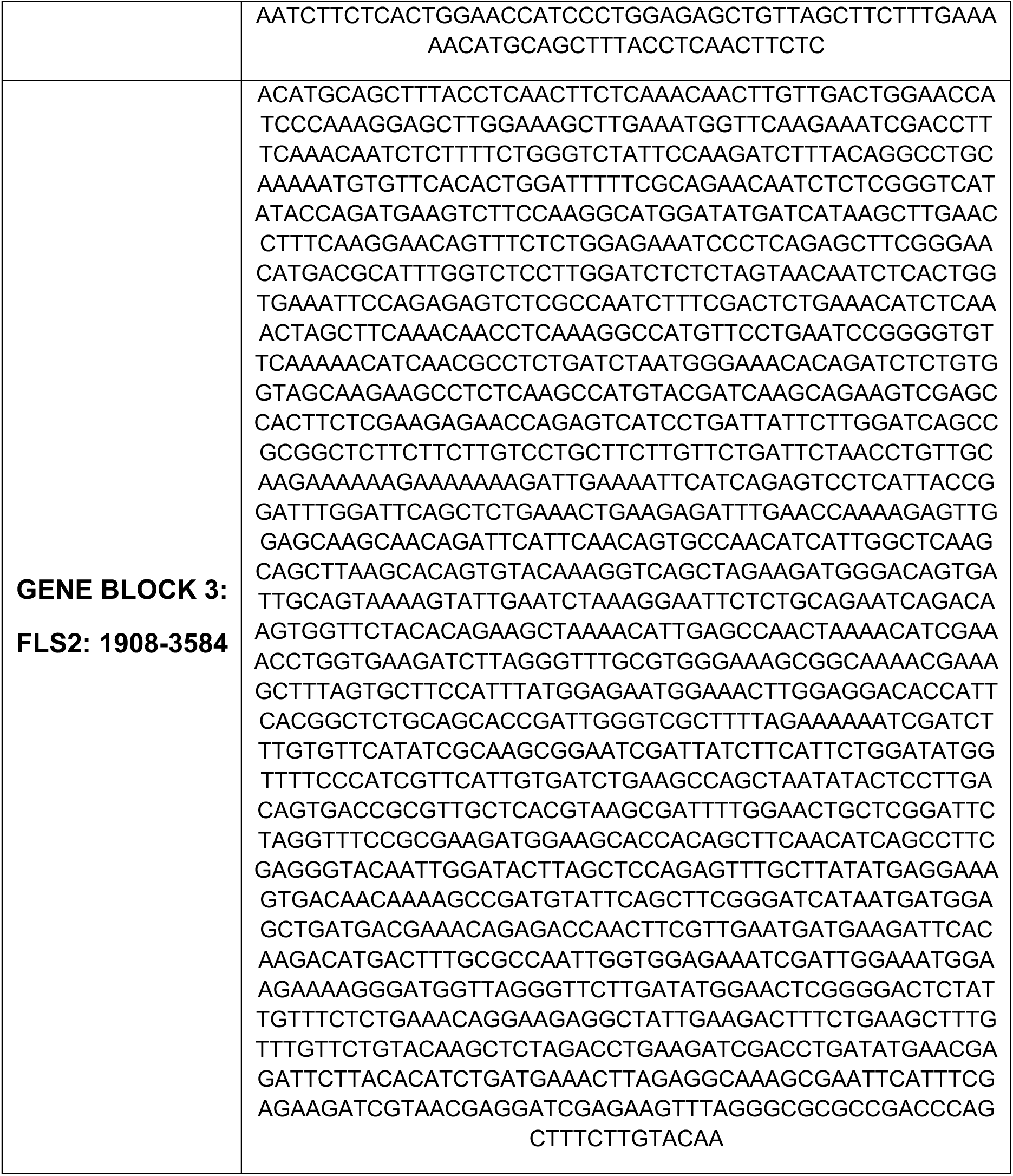
Synthetic gene block sequences used for cloning the *FLS2*pMDC7 construct.

**Supplemental Figure 1.**
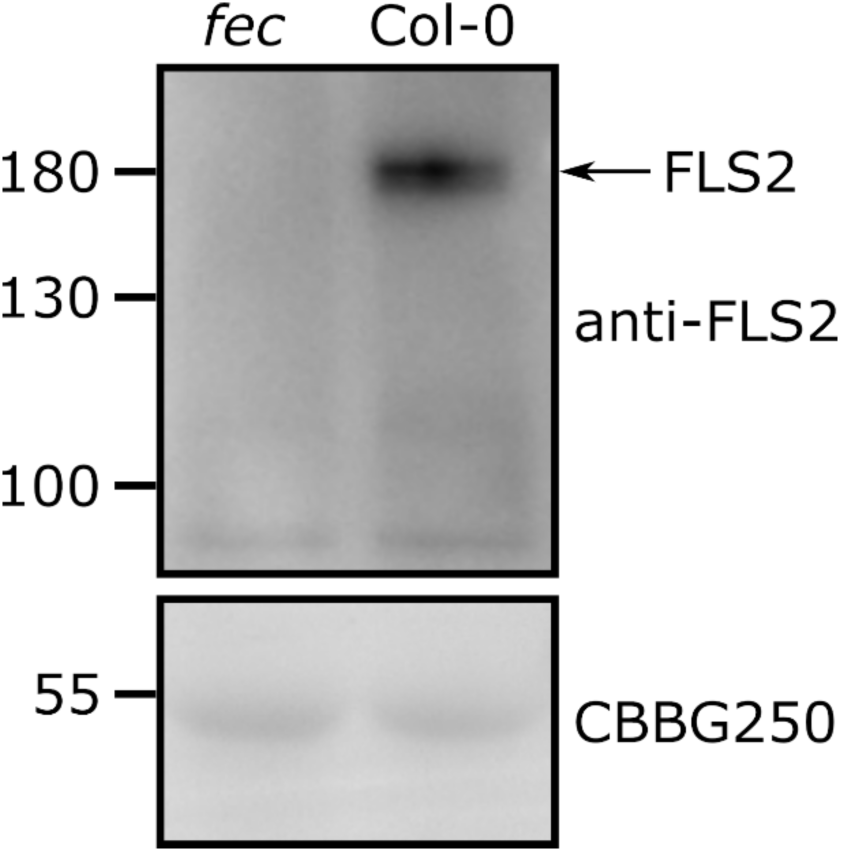
Confirmation of anti-FLS2 antibody specificity. Proteins samples from Col-0 or *fec* seedlings were separated in an 8% SDS-PAGE gel followed by transfer to PVDF and immunoblotting with anti-FLS2 antibodies. After imaging, the membrane was stained with Coomassie Brilliant Blue G250 to show loading.

## REFERENCES

Angelotti F, Hamada E, Bettiol W (2024). A Comprehensive Review of Climate Change and Plant Diseases in Brazil. Plants 13: 2447.

Bita CE, Gerats T (2013). Plant tolerance to high temperature in a changing environment: Scientific fundamentals and production of heat stress-tolerant crops. Frontiers in Plant Science 4: 273.

Bücherl CA, Jarsch IK, Schudoma C, Segonzac C, Mbengue M, Robatzek S, MacLean D, Ott T, Zipfel C (2017). Plant immune and growth receptors share common signalling components but localise to distinct plasma membrane nanodomains. eLife 6: e25114.

Cano-Ramirez DL, Carmona-Salazar L, Morales-Cedillo F, Ramírez-Salcedo J, Cahoon EB, Gavilanes-Ruíz M (2021). Plasma Membrane Fluidity: An Environment Thermal Detector in Plants. Cells 10: 2778.

Carratù L, Franceschelli S, Pardini CL, Kobayashi GS, Horvath I, Vigh L, Maresca B (1996). Membrane lipid perturbation modifies the set point of the temperature of heat shock response in yeast. Proceedings of the National Academy of Sciences of the United States of America 93: 3870.

Caterina MJ, Schumacher MA, Tominaga M, Rosen TA, Levine JD, Julius D (1997). The capsaicin receptor: A heat-activated ion channel in the pain pathway. Nature 389: 816–824.

Chaloner TM, Gurr SJ, Bebber DP (2021). Plant pathogen infection risk tracks global crop yields under climate change. Nature Climate Change 2021 11: 710–715.

Cheng C, Gao X, Feng B, Sheen J, Shan L, He P (2013). Plant immune response to pathogens differs with changing temperatures. Nature Communications 4: 2530.

Cui Y, Li X, Yu M, Li R, Fan L, Zhu Y, Lin J (2018). Sterols regulate endocytic pathways during flg22-induced defense responses in *Arabidopsis*. Development 145.

Curtis MD, Grossniklaus U (2003). A Gateway Cloning Vector Set for High-Throughput Functional Analysis of Genes in Planta. Plant Physiology 133: 462–469.

Desaint H, Aoun N, Deslandes L, Vailleau F, Roux F, Berthomé R (2021). Fight hard or die trying: when plants face pathogens under heat stress. New Phytologist 229: 712– 734.

Deutsch CA, Tewksbury JJ, Huey RB, Sheldon KS, Ghalambor CK, Haak DC, Martin PR (2008). Impacts of climate warming on terrestrial ectotherms across latitude. Proceedings of the National Academy of Sciences of the United States of America 105: 6668–6672.

Felix G, Duran JD, Volko S, Boller T (1999). Plants have a sensitive perception system for the most conserved domain of bacterial flagellin. Plant Journal 18: 265–276.

Finka A, Cuendet AFH, Maathuis FJM, Saidi Y, Goloubinoff P (2012). Plasma membrane cyclic nucleotide gated calcium channels control land plant thermal sensing and acquired thermotolerance. Plant Cell 24: 3333–3348.

Gao F, Han X, Wu J, Zheng S, Shang Z, Sun D, Zhou R, Li B (2012). A heat-activated calcium-permeable channel - Arabidopsis cyclic nucleotide-gated ion channel 6 - Is involved in heat shock responses. Plant Journal 70: 1056–1069.

Garrett KA, Dendy SP, Frank EE, Rouse MN, Travers SE (2006). Climate change effects on plant disease: Genomes to ecosystems. Annual Review of Phytopathology 44: 489–509.

Gómez-Gómez L, Boller T (2000). FLS2: An LRR Receptor–like Kinase Involved in the Perception of the Bacterial Elicitor Flagellin in Arabidopsis. Molecular Cell 5: 1003–1011.

Gronnier J, Franck CM, Stegmann M, Defalco TA, Abarca A, von Arx M, Dünser K, Lin W, Yang Z, Kleine-Vehn J, et al. (2022). Regulation of immune receptor kinase plasma membrane nanoscale organization by a plant peptide hormone and its receptors. eLife 11: e74162.

Hashimoto K, Kudla J (2011). Calcium decoding mechanisms in plants. Biochimie 93: 2054–2059.

Huot B, Castroverde CDM, Velásquez AC, Hubbard E, Pulman JA, Yao J, Childs KL, Tsuda K, Montgomery BL, He SY (2017). Dual impact of elevated temperature on plant defence and bacterial virulence in *Arabidopsis*. Nature Communications 8: 1808.

Hurst CH, Turnbull D, Xhelilaj K, Myles S, Pflughaupt RL, Kopischke M, Davies P, Jones S, Robatzek S, Zipfel C, et al. (2023). S-acylation stabilizes ligand-induced receptor kinase complex formation during plant pattern-triggered immune signaling. Current Biology 33: 1588–1596.e6.

Janda M, Lamparová L, Zubíková A, Burketová L, Martinec J, Krčková Z (2019). Temporary heat stress suppresses PAMP-triggered immunity and resistance to bacteria in Arabidopsis thaliana. Molecular Plant Pathology 20: 1005–1012.

Jeworutzki E, Roelfsema MRG, Anschütz U, Krol E, Elzenga JTM, Felix G, Boller T, Hedrich R, Becker D (2010). Early signaling through the arabidopsis pattern recognition receptors FLS2 and EFR involves Ca^2+^-associated opening of plasma membrane anion channels. Plant Journal 62: 367–378.

Jones JDG, Dangl JL (2006). The plant immune system. Nature 444: 323–329.

Jung JH, Seo PJ, Oh E, Kim J (2023). Temperature perception by plants. Trends in Plant Science 28: 924–940.

Kadota Y, Sklenar J, Derbyshire P, Stransfeld L, Asai S, Ntoukakis V, Jones JD, Shirasu K, Menke F, Jones A, et al. (2014). Direct Regulation of the NADPH Oxidase RBOHD by the PRR-Associated Kinase BIK1 during Plant Immunity. Molecular Cell 54: 43–55.

Kim JH, Castroverde CDM, Huang S, Li C, Hilleary R, Seroka A, Sohrabi R, Medina-Yerena D, Huot B, Wang J, et al. (2022). Increasing the resilience of plant immunity to a warming climate. Nature 607: 339–344.

Knight MR, Campbell AK, Smith SM, Trewavas AJ (1991). Transgenic plant aequorin reports the effects of touch and cold-shock and elicitors on cytoplasmic calcium. Nature 352: 524–526.

Knight H, Mugford SG, Ülker B, Gao D, Thorlby G, Knight MR (2009). Identification of SFR6, a key component in cold acclimation acting post-translationally on CBF function. Plant Journal 58: 97–108.

Kudla J, Becker D, Grill E, Hedrich R, Hippler M, Kummer U, Parniske M, Romeis T, Schumacher K (2018). Advances and current challenges in calcium signaling. New Phytologist 218: 414–431.

Lenzoni G, Liu J, Knight MR (2018). Predicting plant immunity gene expression by identifying the decoding mechanism of calcium signatures. New Phytologist 217: 1598– 1609.

Liu J, Feng L, Li J, He Z (2015). Genetic and epigenetic control of plant heat responses. Frontiers in Plant Science 6: 133426.

Livak KJ, Schmittgen TD (2001). Analysis of relative gene expression data using real-time quantitative PCR and the 2-ΔΔ*C_t_* method. Methods 25: 402–408.

Martinière A, Shvedunova M, Thomson AJW, Evans NH, Penfield S, Runions J, Mcwatters HG (2011). Homeostasis of plasma membrane viscosity in fluctuating temperatures. New Phytologist 192: 328–337.

McAinsh MR, Hetherington AM (1998). Encoding specificity in Ca^2+^ signalling systems. Trends in Plant Science 3: 32–36.

McAinsh MR, Pittman JK (2009). Shaping the calcium signature. New Phytologist 181: 275–294.

Murakami Y, Tsuyama M, Kobayashi Y, Kodama H, Iba K (2000). Trienoic fatty acids and plant tolerance of high temperature. Science 287: 476–479.

Niu Y, Xiang Y (2018). An Overview of Biomembrane Functions in Plant Responses to High-Temperature Stress. Frontiers in Plant Science 9: 915.

Örvar BL, Sangwan V, Omann F, Dhindsa RS (2000). Early steps in cold sensing by plant cells: The role of actin cytoskeleton and membrane fluidity. Plant Journal 23: 785– 794.

Pedersen JZ, Cox RP (1984). Relationship between Thylakoid Membrane Fluidity and the Kinetics of Salt Induced Fluorescence Changes: a Spin Label Study. Advances in Photosynthesis Research: 51–54.

Penfield S (2008). Temperature perception and signal transduction in plants. New Phytologist 179: 615–628.

Poovaiah BW, Du L (2018). Calcium signaling: decoding mechanism of calcium signatures. New Phytologist 217: 1394–1396.

Ranf S, Eschen-Lippold L, Pecher P, Lee J, Scheel D (2011). Interplay between calcium signalling and early signalling elements during defence responses to microbe- or damage-associated molecular patterns. Plant Journal 68: 100–113.

Ranf S, Grimmer J, Pöschl Y, Pecher P, Chinchilla D, Scheel D, Lee J (2012). Defense-related calcium signaling mutants uncovered via a quantitative high-throughput screen in *Arabidopsis thaliana*. Molecular Plant 5: 115–130.

Rasmussen S, Barah P, Suarez-Rodriguez MC, Bressendorff S, Friis P, Costantino P, Bones AM, Nielsen HB, Mundy J (2013). Transcriptome Responses to Combinations of Stresses in Arabidopsis. Plant Physiology 161: 1783–1794.

Saidi Y, Finka A, Goloubinoff P (2011). Heat perception and signalling in plants: A tortuous path to thermotolerance. New Phytologist 190: 556–565.

Saidi Y, Finka A, Muriset M, Bromberg Z, Weiss YG, Maathuis FJM, Goloubinoff P (2009). The heat shock response in moss plants is regulated by specific calcium-permeable channels in the plasma membrane. Plant Cell 21: 2829–2843.

Saidi Y, Peter M, Fink A, Cicekli C, Vigh L, Goloubinoff P (2010). Membrane lipid composition affects plant heat sensing and modulates Ca^2+^-dependent heat shock response. Plant Signaling & Behavior 5: 1530–1533.

Sangwan V, Foulds I, Singh J, Dhindsa RS (2001). Cold-activation of *Brassica napus* BN115 promoter is mediated by structural changes in membranes and cytoskeleton, and requires Ca^2+^ influx. Plant Journal 27: 1–12.

Schroda M, Hemme D, Mühlhaus T (2015). The *Chlamydomonas* heat stress response. Plant Journal 82: 466–480.

Sedaghatmehr M, Thirumalaikumar VP, Kamranfar I, Marmagne A, Masclaux-Daubresse C, Balazadeh S (2019). A regulatory role of autophagy for resetting the memory of heat stress in plants. Plant Cell and Environment 42: 1054–1064.

Shields A, Yao L, Rossi CAM, Collado Cordon P, Kim JH, AlTemen WMA, Li S, Marchetta EJR, Shivnauth V, Chen T, et al. (2025). Warm temperature suppresses plant systemic acquired resistance by intercepting N-hydroxypipecolic acid biosynthesis. The Plant Journal 123: e70374.

Singh BK, Delgado-Baquerizo M, Egidi E, Guirado E, Leach JE, Liu H, Trivedi P. (2023). Climate change impacts on plant pathogens, food security and paths forward. Nature Reviews Microbiology 21: 640–656.

Suzuki N, Rivero RM, Shulaev V, Blumwald E, Mittler R (2014). Abiotic and biotic stress combinations. New Phytologist 203: 32–43.

Tran TM, Ma Z, Triebl A, Nath S, Cheng Y, Gong BQ, Han X, Wang J, Li JF, Wenk MR, et al. (2020). The bacterial quorum sensing signal DSF hijacks *Arabidopsis thaliana* sterol biosynthesis to suppress plant innate immunity. Life science alliance 3.

Vaumourin E, Laine AL (2018). Role of temperature and coinfection in mediating pathogen life-history traits. Frontiers in Plant Science 871: 408482.

Velásquez AC, Castroverde CDM, He SY (2018). Plant–Pathogen Warfare under Changing Climate Conditions. Current Biology 28: R619–R634.

Vilar B, Tan CH, McNaughton PA (2020). Heat detection by the TRPM2 ion channel. Nature 584: E5–E12.

Wang Y, Bao Z, Zhu Y, Hua J (2009a). Analysis of temperature modulation of plant defense against biotrophic microbes. Molecular Plant-Microbe Interactions 22: 498–506.

Wang L, Tsuda K, Sato M, Cohen JD, Katagiri F, Glazebrook J (2009b). Arabidopsis CaM Binding Protein CBP60g Contributes to MAMP-Induced SA Accumulation and Is Involved in Disease Resistance against *Pseudomonas syringae*. PLOS Pathogens 5: e1000301.

Xu H, Ramsey IS, Kotecha SA, Moran MM, Chong JA, Lawson D, Ge P, Lilly J, Silos-Santiago I, Xie Y, et al. (2002). TRPV3 is a calcium-permeable temperature-sensitive cation channel. Nature 418: 181–186.

Yang F, Kimberlin AN, Elowsky CG, Liu Y, Gonzalez-Solis A, Cahoon EB, Alfano JR (2019). A Plant Immune Receptor Degraded by Selective Autophagy. Molecular Plant 12: 113–123.

Zhang Y, Xu S, Ding P, Wang D, Cheng YT, He J, Gao M, Xu F, Li Y, Zhu Z, et al. (2010). Control of salicylic acid synthesis and systemic acquired resistance by two members of a plant-specific family of transcription factors. Proceedings of the National Academy of Sciences of the United States of America 107: 18220–18225.

Zipfel C, Robatzek S, Navarro L, Oakeley EJ, Jones JDG, Felix G, Boller T (2004). Bacterial disease resistance in Arabidopsis through flagellin perception. Nature 428: 764– 767.

